# Enhanced Protection from SARS-CoV-2 Variants by MVA-Based Vaccines Expressing Matched or Mismatched S Proteins Administered Intranasally to hACE2 Mice

**DOI:** 10.1101/2022.12.03.518963

**Authors:** Catherine A. Cotter, Jeffrey L. Americo, Patricia L. Earl, Bernard Moss

## Abstract

The continuous evolution of SARS-CoV-2 strains is contributing to the prolongation of the global pandemic. We previously reported the prevention or more rapid clearance of SARS-CoV-2 from the nasal turbinates and lungs of susceptible K18-hACE2 mice that had been vaccinated intranasally (IN) rather than intramuscularly (IM) with a recombinant MVA (rMVA) expressing a modified S protein of the ancestor SARS-CoV-2 strain. Here, we constructed additional rMVAs and pseudoviruses expressing modified S protein of SARS-CoV-2 variants and compared the ability of vaccines with S proteins that were matched or mismatched to neutralize variants, bind to S proteins and protect K18-hACE2 mice against SARS-CoV-2 challenge. Although vaccines with matched S proteins induced higher neutralizing antibodies, vaccines with mismatched S proteins still protected against severe disease and reduced virus and mRNAs in the lungs and nasal turbinates, though not as well as vaccines with matched S proteins. In mice earlier primed and boosted with rMVA expressing ancestral S, antibodies to the latter increased after one immunization with rMVA expressing Omicron S, but neutralizing antibody to Omicron required a second immunization. Passive transfer of Wuhan immune serum with Omicron S binding but undetectable neutralizing activities reduced infection of the lungs by the variant. Notably, the reduction in infection of the nasal turbinates and lungs was significantly greater when the rMVAs were administered IN rather than IM and this held true for vaccines that were matched or mismatched to the challenge SARS-CoV-2.

## INTRODUCTION

The speed with which safe and efficacious SARS-CoV-2 vaccines were developed was a remarkable achievement. Clinical trials indicated that the mRNA vaccines were 94 to 95% effective in preventing COVID-19 illness ^1,2^ and adenovirus-based vaccines were about 74% effective ^3,4^. Those and most other vaccines are based on the spike (S) protein, which mediates entry of the virus into cells. Initially, it was considered that the proof-reading mechanism employed by coronaviruses would greatly retard the development of escape mutants ^5^. However, successive waves of variants and subvariants appeared with mutations in S including the receptor binding domain (RBD) and some such as Beta and Omicron exhibited resistance to antibodies elicited by ancestor strains ^6^. Nevertheless, boosting with the original vaccines reduce serious disease, though they appear less effective in preventing infection and transmission ^7^. Updated SARS-CoV-2 mRNA vaccines are based on expression of two S proteins: one from an ancestor and the other from a recent isolate ^8^. Another consideration is whether intranasal (IN) or aerosol delivery would prevent infection and transmission better than intramuscular (IM) vaccination.

Recombinant poxvirus platforms are valuable for identifying targets of humoral and cellular immunity, have been developed into numerous veterinary vaccines and are undergoing clinical evaluation for vaccines against unrelated pathogens including SARS-CoV-2 and cancer ^9–11^. We and others described animal studies supporting use of the host-range restricted vaccinia virus Ankara (MVA) as an alternative vector for COVID-19 vaccines ^12–16^. Recent animal studies demonstrated advantages of IN delivery of recombinant MVAs (rMVAs) expressing S ^17–19^. Anti-SARS-CoV-2 IgA and IgG as well as specific T cells were detected in the lungs of IN vaccinated mice and virus was diminished in the upper and lower respiratory tracts following challenge of K18-hACE2 mice with SARS-CoV-2. Here we describe the construction and immunogenicity of rMVAs expressing the S proteins of several variant SARS-CoV-2 strains. The neutralizing and S binding activities of sera following matched and mismatched rMVA boosts were determined as well as protection of K18-hACE2 mice vaccinated IN and IM and challenged IN with SARS-CoV-2 variants. Vaccines that produced low neutralizing activities to mismatched SARS-CoV-2 variants still provided durable protection, but vaccines matched to the challenge virus elicited higher neutralizing activities and were more effective. For both matched and mismatched immunizations, the IN route was better than IM at reducing virus infection of the upper and lower respiratory tracts. In mice earlier primed and boosted with rMVA expressing ancestral S, antibodies to the latter increased after one immunization with rMVA expressing Omicron S, but neutralizing antibody to Omicron required a second immunization.

## RESULTS

### Relative virulence of SAR-CoV-2 variants in the K18-hACE2 mouse model

We previously reported that serum from mice vaccinated with an rMVA expressing the spike protein of the ancestor Wuhan strain of SARS-CoV-2, that was modified to stabilize the pre-fusion structure and enhance membrane localization (rMVA-W), neutralized recombinant vesicular stomatitis virus (rVSV) pseudoviruses expressing divergent S proteins to varying degrees ^18^. This cross-reactivity led us to evaluate the ability of rMVA-W and rMVAs expressing variant S proteins to protect K18-hACE2 mice against challenge with SARS-CoV-2 variants. Before undertaking protection studies, we compared the relative virulence of four SARS-CoV-2 variants, CoV-Washington (W, S identical to Wuhan), -Beta (B), -Delta (D), and -Omicron (O, BA.1.1) in the mouse model system. Following IN infection of the hACE2 mice with CoV-W, -B and -D, weight loss was detected with a dose of 10^2^ TCID_50_ (Fig. S1A–C) and 100% death occurred with 10^4^ TCID50 (Fig. S1D-F). CoV-O was less virulent than the others, with only two of five mice succumbing on day 7 to a dose of 5×10^4^ (Fig. S1G). For a further comparison, the hACE2 mice were inoculated IN with varying doses of CoV-W or -O and the amounts of virus in the lungs and nasal turbinates determined. The lungs of CoV-W-infected mice contained 100-to 1,000-fold more virus than the lungs of CoV-O-inoculated mice on day 2 (Fig. S1H). At that early time, virus was detected in the nasal turbinates only of mice inoculated with CoV-W (Fig. S1I). Our finding that CoV-O is less virulent than other variants in mice is in accord with other studies ^20,21^. In subsequent challenge experiments, a dose of 5×10^4^ TCID_50_ (highest possible dose due to titer) was used for CoV-O and 10^4^ TCID_50_ for the others.

### Protective immunity to SARS-CoV-2 variants following IM vaccination with rMVA-W

K18-hACE2 mice were vaccinated IM twice 3 weeks apart with the parental MVA as a control or with rMVA-W and challenged with CoV variants 2 weeks later (Fig. 1A). Antibody binding to the Wuhan RBD was detected by ELISA after the first immunization and was not increased significantly by the second (Fig. 1B). Pseudoviruses, named for the spike protein that they express, were neutralized in the order rVSV-W > D > -B after the first immunization (Fig. 1C). The neutralization titers increased significantly after the second immunization for the vaccine mismatched rVSV-B and rVSV-D but not for the matched rVSV-W, which was already high.

**Fig. 1.**
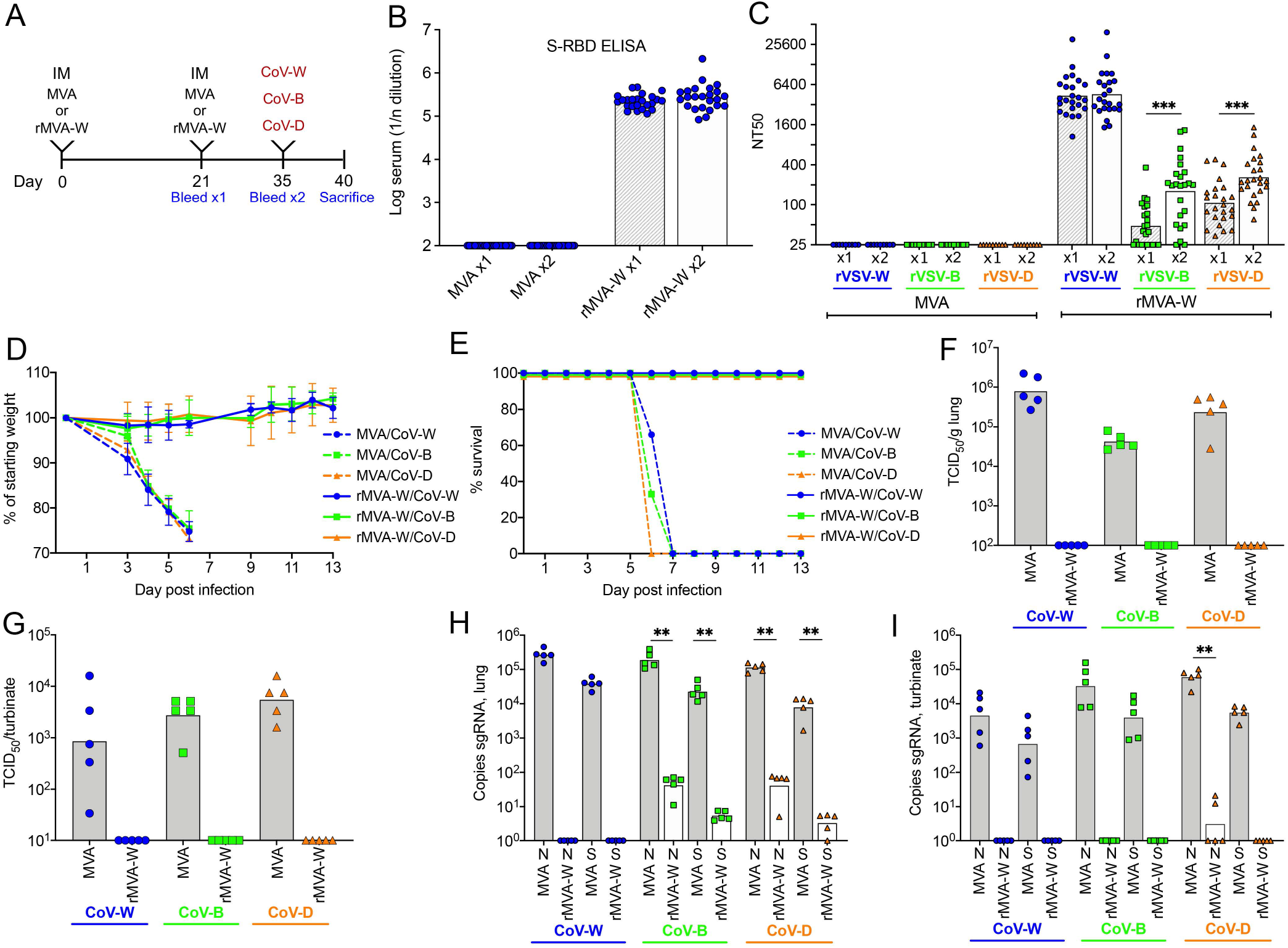
Protective immunity to SARS-CoV-2 variants following IM vaccination with rMVA-W. (**A**) K18-hACE2 mice were vaccinated IM twice with MVA control (n=40) or rMVA-W (n=40), divided into groups of n=10 and challenged with CoV-W, -B, or -D two weeks later. (**B, C**) Serum antibody binding to the Wuhan S RBD and neutralization of pseudoviruses rVSV-W, - B or -D at 3 and 5 weeks after vaccination. (**D, E**) Time course of weight loss and survival on days after challenge. (**F, G**) Recovery of SARS-CoV-2 from lungs and nasal turbinates on day 5 after challenge. (**H, I**) Copies of sgRNAs N and S normalized to 18s RNA from lungs and nasal turbinates on day 5 after challenge. Abbreviation: D, day; sac, sacrifice. * p=/< 0.03; ** p=/<0.002; *** p=/<0.0002; **** p<0.0001. Significance not calculated when values of one group were all below the limit of detection.

Following challenge with CoV-W, -B or -D, mice that received the control MVA vector lost weight and died by day 6, whereas mice that received rMVA-W lost no weight and survived infection with each of the variants (Fig. 1D, E). Five additional mice in each group were sacrificed on day 5, prior to severe morbidity, for analysis of virus titers and subgenomic (sg) RNAs in internal organs. Substantial amounts of each of the SARS-CoV-2 viruses were detected by infectivity assays in the lungs and turbinates of control mice that received the MVA vector, whereas none was detected in mice that received rMVA-W regardless of the challenge strain of virus (Fig. 1F, G). Analysis of sgRNAs provides an alternative and more sensitive assay for replication than the titer of infectious SARS-CoV-2 from tissues ^22^ and sgN is most abundant followed by sgS ^23^. Digital droplet polymerase chain reaction (ddPCR) was used to detect sgN and sgS in our study and the values normalized to 18s ribosomal RNA in each sample. High levels of sgN and sgS RNAs were detected in the lungs of the control mice that received the MVA vector following challenge with each of the variants. In contrast, sgRNAs were not detected in mice immunized with rMVA-W and challenged with CoV-W and the levels were significantly diminished in mice challenged with CoV-B and -D (Fig. 1H). High titers of sgRNAs were also present in the nasal turbinates of challenged mice that had been immunized with the control MVA but were undetected or significantly diminished in mice that had been immunized with rMVA-W (Fig. 1I). These results indicated that a vaccine expressing the Wuhan S and administered IM protected against weight loss and death and significantly reduced the replication of CoV-B and -D by day 5, despite lower levels of neutralizing antibody to their S proteins. Nevertheless, virus replication in the lungs as determined by sgRNAs was lowest when the S proteins of the vaccine and challenge were matched.

### Duration of cross-protective immunity following IM vaccination with rMVA-W

In the above experiment, mice were challenged at 3 weeks after vaccination when antibody levels were near their peak. To determine the duration of cross-protective immunity, additional groups of K18-hACE2 mice were vaccinated IM twice and held for 9 months (Fig. 2A). Over this period, the binding to the Wuhan S protein decreased by about 70% (Fig. 2B). The neutralizing titer for each variant was boosted by the second immunization but then each also decreased substantially over time (Fig. 2C).

**Fig. 2.**
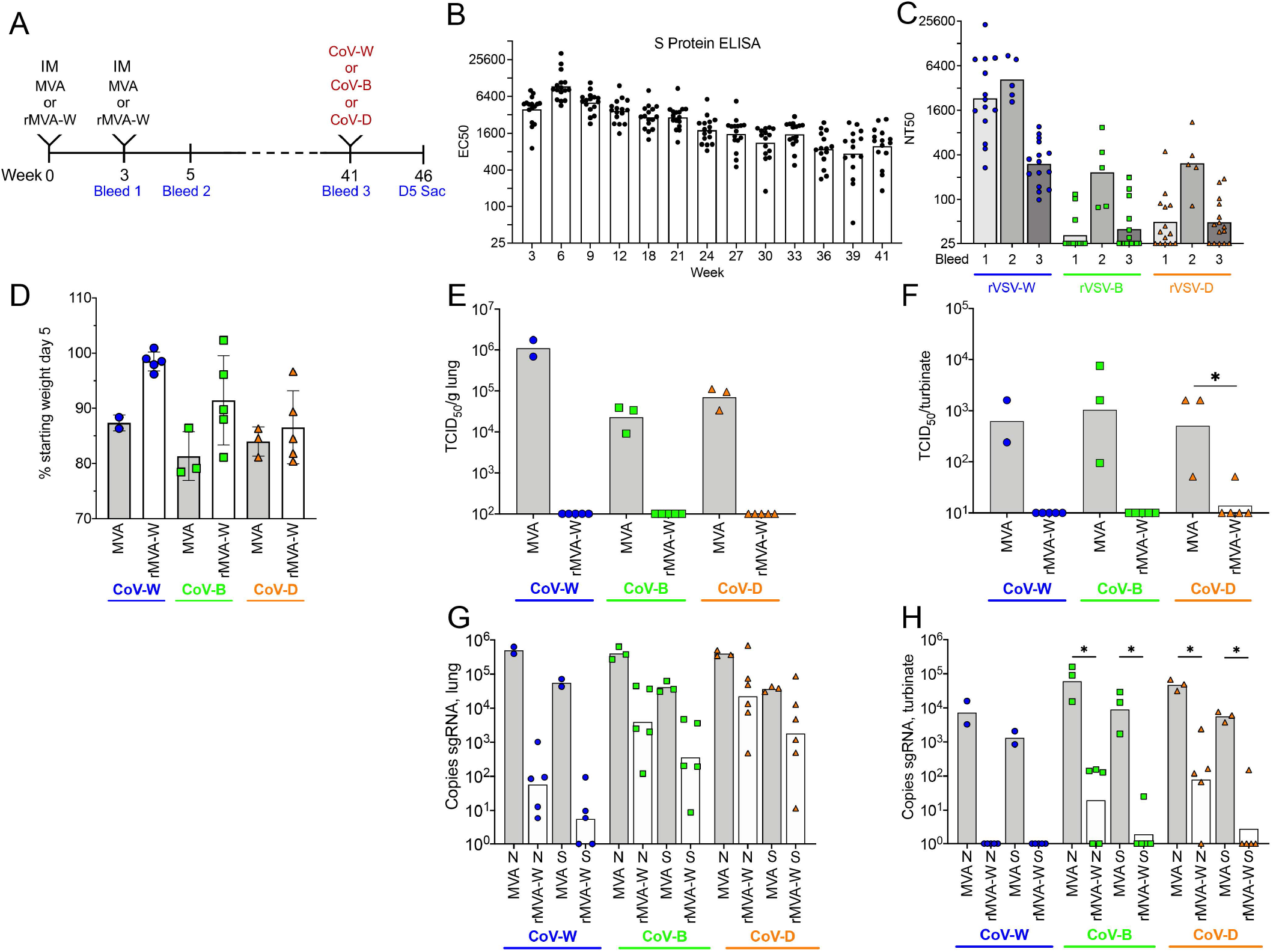
Duration of cross-protective immunity following IM vaccination with rMVA-W. (**A**) K18-hACE2 mice were vaccinated IM twice with MVA control (n=8) divided into groups of 2 – 3 or rMVA-W (n=15), divided into groups 5 and challenged with CoV-W, -B, or -D approximately 9 months later. (**B**) Binding of serum antibodies to Wuhan S protein determined by ELISA. (**C**) Neutralization of pseudoviruses rVSV-W, -B and -D by serum obtained at weeks 3, 5 and 41. (**D**) Weights of mice on day 5 after challenge relative to starting weights. (**E, F**) Recovery of SARS-CoV-2 from lungs and nasal turbinates on day 5 after challenge. (**G, H**) Copies of sgRNAs N and S normalized to 18s RNA from lungs and nasal turbinates on day 5 after challenge.

After challenge with the SARS-CoV-2 variants, mice that had received the control MVA vector lost ~10 to 20% of their starting weight by day 5; mice immunized with rMVA-W and challenged with CoV-W lost little weight while those challenged with CoV-B and -D were intermediate (Fig. 2D). Mice that received the control MVA vector had substantial amounts of virus in the lungs and turbinates on day 5 following challenge, whereas mice that had been vaccinated with rMVA-W had little or no virus regardless of which SARS-CoV-2 strain was used for challenge (Fig. 2E, F). Substantial differences, however, were revealed by analysis of sgRNAs. Mice vaccinated with rMVA-W and challenged with CoV-W had >3-log mean reduction of sgRNAs in the lungs compared to controls whereas mice challenged with CoV-B and -D had 1- to 2-log mean reductions (Fig. 2G). The same trend was observed in the nasal turbinates: mice challenged with CoV-W had no detectable sgRNAs, whereas sgRNAs were significantly reduced relative to controls in mice challenged with CoV-B and -D (Fig. 2H). We concluded that protection was durable, but at a reduced level at 9 months and was greater when the vaccine and challenge virus were matched.

### Comparison of IM and IN immunizations with rMVA-W on inhibition of early stages of variant infections

The experiments described thus far demonstrated that IM vaccination with rMVA-W protected mice from lethal infection with Beta and Delta strains and reduced virus and sgRNAs present in the lungs and nasal turbinates by day 5 after challenge. For the next experiments, we analyzed virus titers and sgRNAs at 2 and 4 days after infection with SARS-CoV-2 to determine the effect of vaccination on earlier stages of infection (Fig. 3A). Binding to the Wuhan S RBD was detected after the first immunization and increased slightly after the second (Fig. 3B). Neutralization titers obtained with pseudoviruses expressing variant spike proteins were in the order rVSV-W >-D >-B >-O (Fig. 3C). Titers were boosted by the second immunization but, even then, less than half of the mice had serum that neutralized rVSV-O above the limit of detection.

**Fig. 3.**
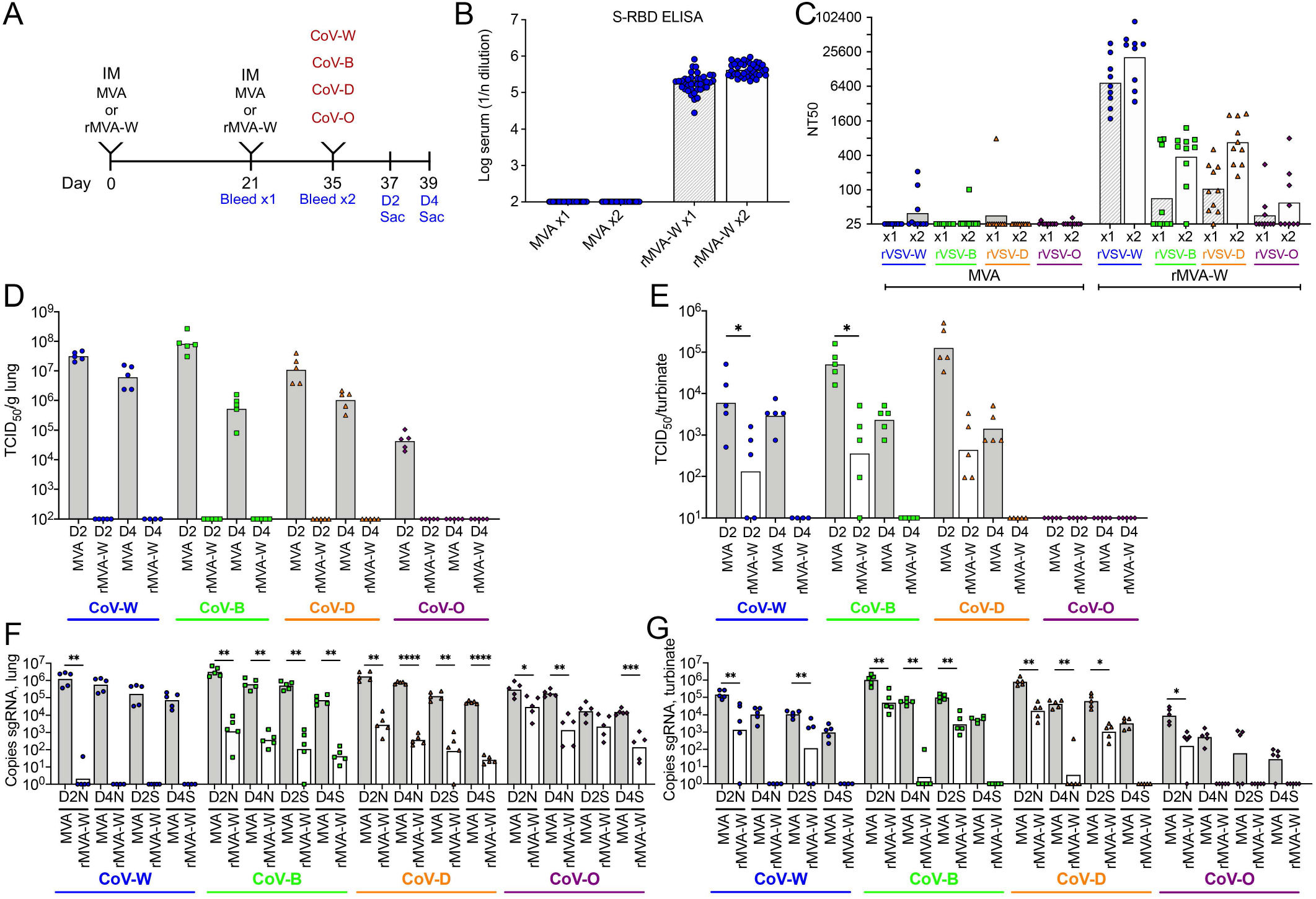
Inhibition of early stages of variant SARS-CoV-2 infections in mice immunized IM with rMVA-W. (**A**) K18-hACE2 mice were vaccinated IM twice with MVA control (n=40) or rMVA-W (n=40), divided into groups of 10 and challenged 2 weeks later with CoV-W, -B, -D or -O 3. (**B**) Binding of serum antibodies to Wuhan RBD after first and second immunizations determined by ELISA. (**C**) Neutralization of pseudoviruses rVSV-W, -B, -D and -O by sera obtained after the first and second immunizations. (**D, E**) Recovery of SARS-CoV-2 from lungs and nasal turbinates on days 2 and 4 after challenge. (**F, G)**Copies of sgRNAs N and S normalized to 18s RNA from lungs and nasal turbinates on days 2 and 4 after challenge.

Following challenge of mice receiving the control MVA vector, the virus titers in the lungs were highest on day 2 and reduced to varying degrees on day 4 for each variant (Fig. 3D). Notably, no virus was recovered from the lungs on either day from mice vaccinated with rMVA-W and challenged with CoV-W, -B, -D or -O. Mice that received the control MVA vector and were subsequently challenged had substantial amounts of virus in the nasal turbinates that were higher on day 2 than day 4 except for CoV-O, in which none was detected (Fig. 3E). For mice vaccinated with rMVA-W, the CoV-W, -B or -D titers in the turbinates were reduced relative to the controls on day 2 and undetectable on day 4. The presence of SARS-CoV-2 in the turbinates of vaccinated mice was missed in the previous experiment (Fig. 2F) when only day 5 was examined. High amounts of sgRNAs were found in the lungs of mice inoculated with the control MVA, on days 2 and 4 after challenge with the variants including CoV-O (Fig. 3F). In contrast to the controls, mice that had been immunized with rMVA-W and challenged with CoV-W had little or no sgRNA in the lungs on either day, whereas virus was reduced but still detected on both days after challenge with the other variants (Fig. 3F). In the nasal turbinates, sgRNAs were detected on days 2 and 4 of mice that received the control MVA vector and challenged with each of the variants including CoV-O, whereas sgRNAs were detected in vaccinated mice challenged with CoV-W or variants only on day 2 (Fig. 3G). These data indicated that IM vaccination with rMVA-W reduced replication of each of the variants but did not prevent infection as judged by the detection of virus and sgRNAs in the lungs and turbinates. Interestingly, although rMVA-W induced low Omicron neutralizing activity as measured *in vitro*, the lung titers of Omicron were significantly reduced in vaccinated mice.

We previously reported that IN administration of rMVA-W prevented or more rapidly eliminated upper respiratory infection with CoV-W than when administered IM ^18^. Here, we wanted to determine whether IN delivery is advantageous for cross-protection of variants. Mice were inoculated with the MVA control or rMVA-W as in the previous experiment, except that the route was IN (Fig. 4A). Antibody binding to the RBD of the Wuhan spike protein was detected after the first immunization with rMVA-W and increased slightly after the second (Fig. 4B). The second immunization with rMVA-W significantly increased neutralizing antibodies in the blood to each of the variants except Omicron (Fig. 4C), as was the case for IM immunization. Following challenge of mice that received the control MVA, the CoV-W and variant strains were detected in the lungs and turbinates on day 2 and decreased on day 4 (Fig. 4D, E). In addition, considerable amounts of CoV-B and lesser amounts of CoV-D were recovered from the brains of the control mice on day 4 (Fig. 4F). However, in mice vaccinated IN with rMVA-W, virtually no virus of any strain was detected on days 2 or 4 in lungs, nasal turbinates or brains of mice (Fig. 4D-F), whereas considerable virus had been detected on day 2 in the turbinates of mice that had been vaccinated IM (Fig. 3E). Furthermore, sgRNAs were greatly reduced on day 2 and virtually absent on day 4 in the lungs of IN-vaccinated mice following challenge with CoV-W, -B or -D and were significantly reduced after challenge with CoV-O compared to controls (Fig. 4G). Moreover, sgRNAs were undetectable or detected at very low levels in the turbinates of mice infected with any of the variants on day 2 and none was detected on day 4 (Fig. 4H), in contrast to the results obtained by IM vaccination (Fig. 3G). For each variant, there was a greater reduction of sgRNAs in mice immunized IN compared to IM as depicted for the nasal turbinates on day 2 (Fig. 4I). Thus, IN vaccination with rMVA-W was advantageous for protection against variant as well as matched challenges.

**Fig. 4.**
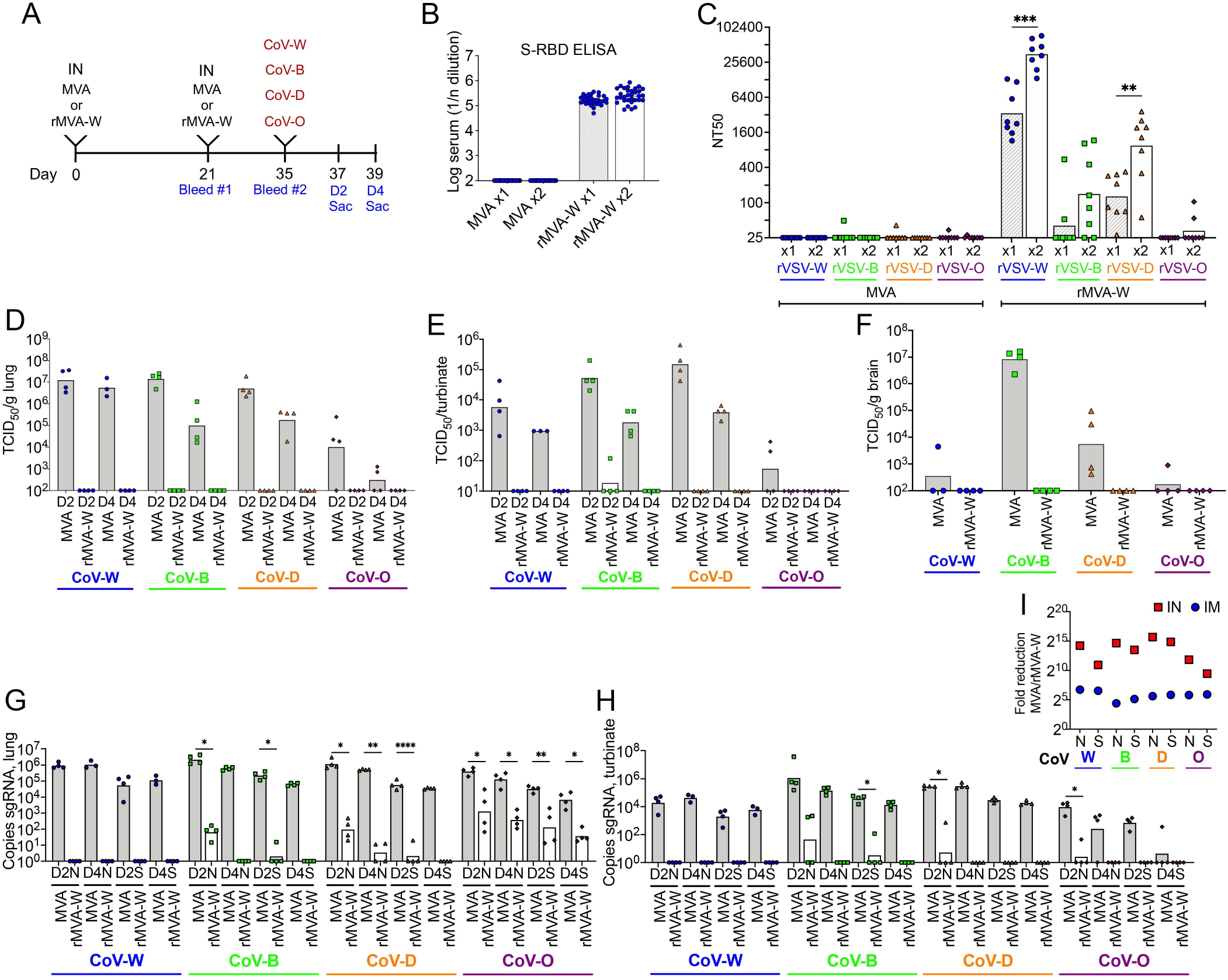
Greater inhibition of early stages of variant SARS-CoV-2 infections in mice immunized IN with rMVA-W. (**A**) K18-hACE2 mice were vaccinated IN twice with MVA control (n=31) or rMVA-W (n=32), divided into groups of 7 or 8 and 2 weeks later challenged with CoV-W, -B, or -D. (**B**) Serum antibody binding to Wuhan RBD determined by ELISA after the first and second immunizations. (**C**) Neutralization of pseudoviruses rVSV-W, -B, -D and -O by serum obtained after the first and second immunizations. (**D, E, F**) Recovery of SARS-CoV-2 from lungs and nasal turbinates on days 2 and brain on day 4 after challenge. (**G,H**) Copies of sgRNAs N and S normalized to 18s RNA from lungs and nasal turbinates on days 2 and 4 after challenge. (**I**) Fold-reduction of sgRNAs in nasal turbinates on day 2 for mice immunized IN and IM. Data replotted from Fig. 3G and 4H.

### Construction and immunogenicity of rMVAs with variant spikes

Thus far, we immunized mice with rMVA expressing the Wuhan S protein and challenged with variant SARS-CoV-2. For the next experiments, we constructed rMVAs expressing the S proteins of variant strains. Equivalent S protein expression was verified by analysis of infected HeLa cells by Western blotting using antibody to the FLAG tag (Fig. S2A, B), although there was some difference in binding of anti-Wuhan RBD to the variants reflecting sequence differences (Fig. S2A, C). Cell surface expression of the S proteins and binding to hACE2 were demonstrated by flow cytometry (Fig. S2D). To compare their immunogenicity, C57BL/6 mice were vaccinated IM with rMVAs expressing variant S proteins followed by a boost with the same rMVA used for the first vaccination (Fig. 5A). The neutralization of pseudoviruses expressing spike proteins matched or mismatched to the vaccines were determined (Fig. 5B). Sera from mice immunized with rMVA-W neutralized rVSV-W significantly better than rVSV-D and had no detectable activity against rVSV-O. After the second vaccination with rMVA-W, neutralizing antibody to rVSV-W and -D were increased but most of the mice still made no detectable neutralizing antibody to rVSV-O. Sera from mice vaccinated with rMVA-D neutralized rVSV-W almost as well as rVSV-D but also had low neutralizing activity to rVSV-O and only small increases occurred after the second immunization. Sera from mice that received rMVA-O had little or no neutralizing activity to rVSV-W or -D but had detectable activity against the matching rVSV-O. After two and three rMVA-O vaccinations, neutralization of rVSV-O was boosted but the neutralizing titers to rVSV-W and -D were minimally increased.

**Fig. 5.**
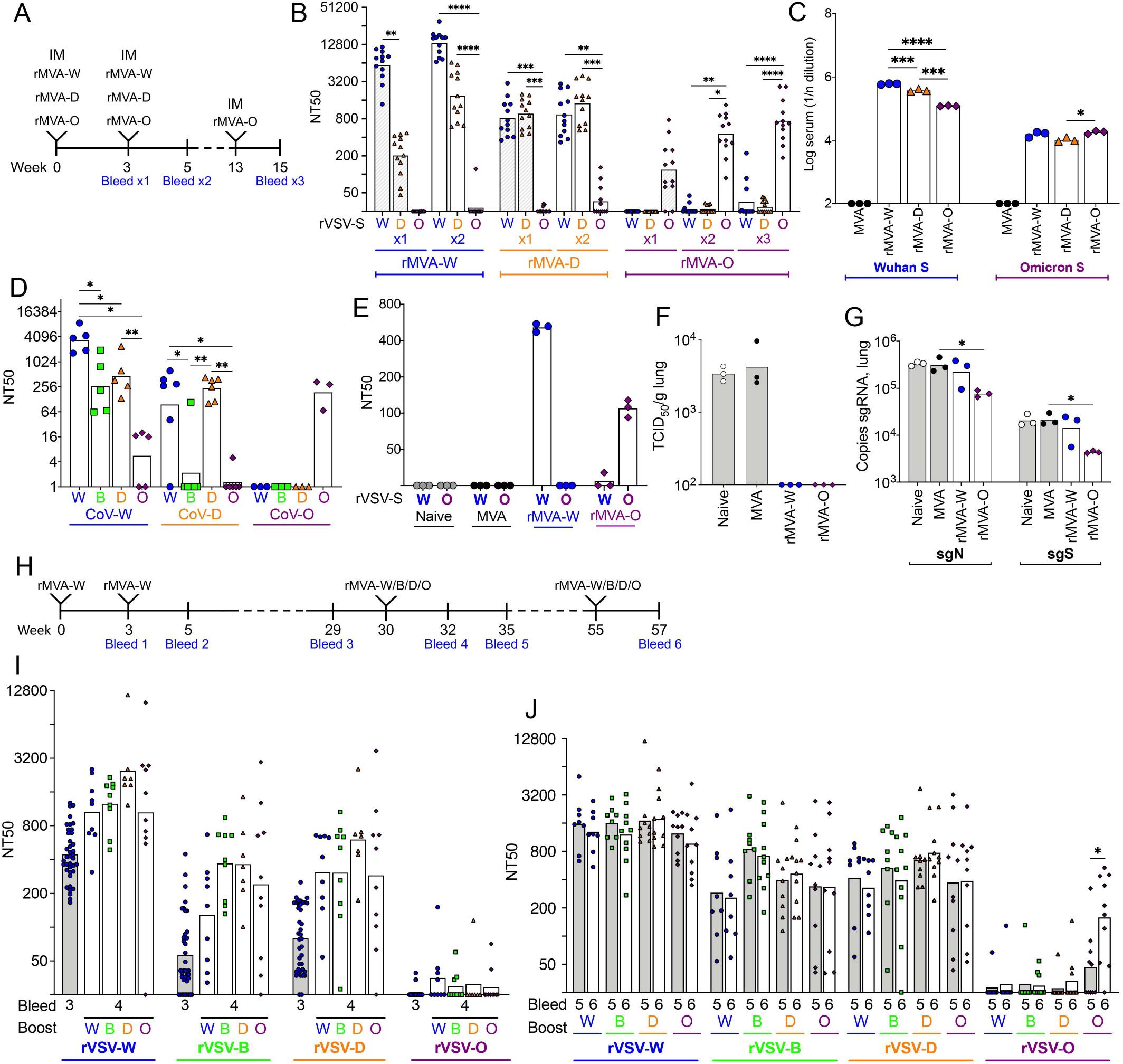
Neutralizing antibody responses following boosts with rMVAs expressing matched or mismatched S. (**A-C, E-**) Matched prime and boost IM vaccinations of C57BL/6 mice (n= 40 per group) with rMVA-W, -D, and -O. (A) Timeline. (B) Neutralization of pseudoviruses rVSV-W, -D and -O by serum obtained after the prime with rMVA-W, -D and -O and after the matched boost. (**C**) Binding of pooled serum from the second bleed to Wuhan and Omicron S determined by ELISA. (D) Neutralization of pseudoviruses by sera from mice sublethally infected with CoV-W, CoV-D or CoV-O, (**E**) Serum neutralizing titers of mice one day after receiving pooled serum IP from naïve mice or mice immunized twice with MVA, rMVA-W or rMVA-O. (**F, G**) Recovery of SARS-CoV-2 and sgRNAs from the lungs at 4 days after challenge of passively immunized mice. (**H, I**) C57BL/6 mice immunized twice with rMVA-W and boosted with rMVA-W, -B, -D, or -O. (**H**) Timeline. (**I**) Neutralization of pseudoviruses rVSV-W, -B, -D and -O by serum obtained after the third immunization. (i) Neutralization of pseudoviruses by sera obtained after the fourth immunization.

The S proteins expressed by the rMVA vectors had been modified to stabilize the pre-fusion form, prevent furin cleavage and increase cell surface expression as described previously ^12^. To be certain that these changes were not responsible for diminished cross-neutralizing activity, we obtained sera from K18-hACE2 mice that had been sublethally infected with CoV-W, CoV-B, CoV-D or CoV-O in the experiment of Fig. S1). The cross-neutralizing activities of these serum samples were similar to serum obtained from vaccinated mice (Fig. 5D).

In contrast to the low neutralizing antibody, sera from mice immunized twice with rMVA-W exhibited substantial binding to the S proteins of Delta and Omicron, although significantly less than to the Wuhan S protein, whereas sera from mice immunized multiple times with rMVA-O bound to similar extents to Wuhan and Omicron S proteins (Fig. 5C). We considered that the relatively greater binding of sera to mismatched S proteins compared to their neutralizing ability may have significance for cross-protection *in vivo*. To investigate this, pooled sera from mice vaccinated IM with rMVA-W or rMVA-O were inoculated IP into K18-hACE2 mice. One day after serum transfer and just before challenge with CoV-O, the NT50 values of the sera from rMVA-W-vaccinated mice were >400 for rVSV-W and undetectable for rVSV-O (Fig. 5E). The corresponding NT50 values for the mice receiving sera from rMVA-O-vaccinated mice were >100 for rVSV-O and undetectable or barely detectable for rVSV-W. Despite the difference in neutralizing titers, no CoV-O was recovered from the lungs at 4 days after challenge of mice receiving anti-Wuhan or anti-Omicron sera (Fig. 5F). However, the mice that received anti-Omicron serum also had reduced sgN and sgS RNAs in the lungs, whereas the mice that received anti-Wuhan serum did not (Fig. 5G). These data indicated that the anti-Wuhan serum without detectable *in vitro* neutralizing activity for Omicron was partially protective *in vivo* but that the neutralizing anti-Omicron serum was more potent

### Neutralizing antibody responses to mismatched prime and boost vaccinations

A related question is whether heterologous boosting with variant spike proteins will increase neutralizing activity to the original spike protein, the variant or both. To investigate this, C57BL/6 mice were vaccinated twice with rMVA-W. After 30 weeks, the mice were bled and groups of 9 to 10 mice were re-vaccinated with rMVA-W, -B, -D or -O (Fig. 5H). The neutralizing titer against the original immunogen rVSV-W as well as rVSV-B and rVSV-D increased after boosting with each of the variant rMVAs with the exception of rVSV-O (Fig. 5I). Because of the low neutralization titer against Omicron obtained following immunization twice with rMVA-W and once with rMVA-O or other rMVA variants, we decided to boost all the mice a second time with the same rMVAs used in the previous boost. None of the boosts increased the neutralizing titer to rVSV-W, rVSV-B or rVSV-D. However, the rMVA-O boost significantly increased the neutralizing titer to rVSV-O, whereas none of the other boosts did (Fig. 5J). These results indicated that a second immunization with rMVA-O is beneficial both for naïve mice as well as mice primed with the rMVA expressing the ancestor S protein.

### Protective immunogenicity of rMVAs expressing variant S proteins

Next, we analyzed replication of CoV-W and CoV-O in mice that had been vaccinated IM with rMVA-W or rMVA-O (Fig. 6A). As shown earlier in this study, immunization of K18-hACE2 mice with rMVA-W elicited high neutralizing antibody to rVSV-W and very little to rVSV-O, whereas the converse occurred following immunization with rMVA-O (Fig. 6B). Nevertheless, Immunization with either rMVA-W or rMVA-O significantly reduced the titer of CoV-W in the lungs (Fig. 6C) and turbinates (Fig. 6D) although immunization with rMVA-W was more effective on day 2. No virus was detected in the lungs or turbinates of mice immunized with either rMVA-W or rMVA-O and challenged with CoV-O, although it is important to note the low amount of CoV-O in the lungs and barely detectable CoV-O in the nasal turbinates.

**Fig. 6.**
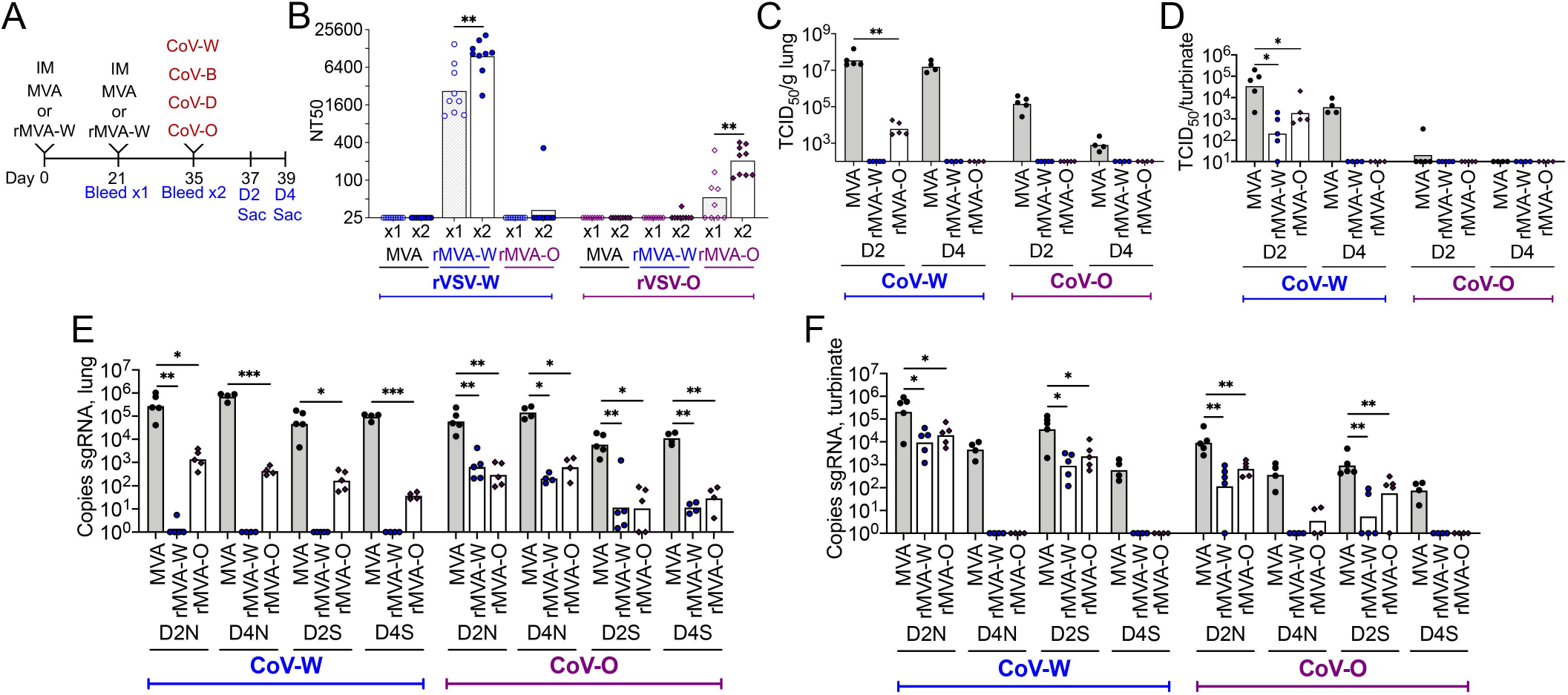
Protection of SARS-CoV-2 challenged mice that received matched or mismatched rMVAs IM. (**A**) Timeline showing IM immunizations of K18-hACE2 mice (n=9 per group) with MVA, rMVA-W and rMVA-O and matched and mismatched challenges with CoV-W and CoV-O. (**B**) Neutralization of pseudoviruses rVSV-W and -O by serum obtained after one and two IM immunizations with MVA, rMVA-W or rMVA-O. (**C, D**) Recovery of SARS-CoV-2 from lungs and nasal turbinates on days 2 and 4 after challenge with CoV-W or CoV-O. (**E, F**) Copies of sgRNAs N and S normalized to 18s RNA from lungs and nasal turbinates on days 2 and 4 after challenge.

Analysis of sgRNAs provided a better basis for comparison of the protection afforded by the different immunizations. Immunization with rMVA-W provided complete protection of the lungs from CoV-W on days 2 and 4, whereas immunization with rMVA-O provided significant but partial protection (Fig. 6E). Nevertheless, the two vaccines provided similar 2-log reduction of sgRNAs in the lungs of mice infected with CoV-O (Fig. 6E). In the nasal turbinates, sgRNAs were reduced on day 2 and undetectable on day 4 after challenge with CoV-W regardless of whether the mice were immunized with rMVA-W or rMVA-O (Fig. 6F). CoV-O sgRNAs were also reduced by similar amounts in the nasal turbinates when vaccinated with either rMVA-W and rMVA-O. These data indicated that significant protection can occur even if low neutralizing antibody is induced.

Next, we investigated the use of rMVA-O as a nasal vaccine (Fig. 7A). After the second immunization with rMVA-O, neutralizing antibody to rVSV-O was similar or higher than that obtained by IM but again there was little or no neutralization of rVSV-W (Fig. 7B). The control mice challenged with CoV-W succumbed to the infection, whereas those challenged with CoV-O had only a transient weight loss due to the low pathogenicity of the latter (Fig. 7C). Mice vaccinated IN with rMVA-O had no weight loss following challenge with either CoV-W or CoV-O (Fig. 7C). There was low or no recovery of virus (Fig. 7D) and significantly diminished sgRNAs in the lungs of mice challenged with either CoV-W or CoV-O (Fig. 7E). Even more striking was the reduction of virus (Fig. 7F) and sgRNAs (Fig. 7G) in the nasal turbinates of mice challenged with CoV-W or CoV-O. The much greater reduction of sgRNAs following IN vaccination than IM vaccination is shown for the turbinates on day 2 in Fig. 7H.

**Fig. 7.**
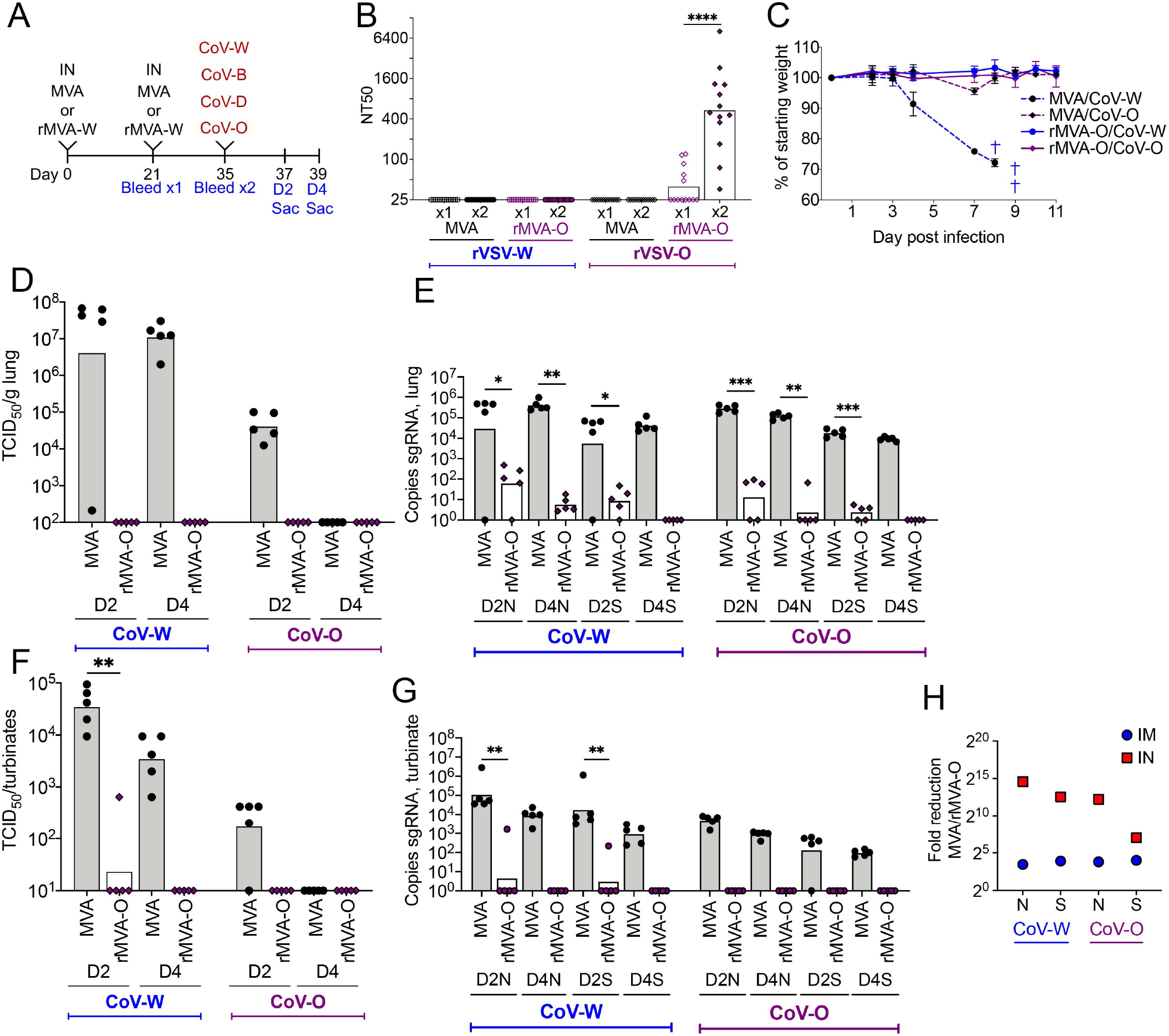
Protection of CoV-W and CoV-O challenged mice vaccinated IN with rMVA-O. (**A**) Timeline of IN immunizations of K18-hACE2 mice (n=13 per group) with MVA or rMVA-O and matched and mismatched challenges with CoV-W and CoV-O. (**B**) Neutralization of pseudoviruses rVSV-W and -O by serum obtained after one and two IN immunizations with MVA or rMVA-O. (**C**) Weight loss of mice following challenge with CoV-O or CoV-W. (D, E) Recovery of virus and sgRNAs in the lungs of mice challenged with CoV-W or CoV-O. (F, G) Recovery of virus and sgRNAs in the nasal turbinates of mice challenged with CoV-W or CoV-O. (H) Fold-reduction of sgRNAs in nasal turbinates on day 2 for mice immunized IN and IM.

## DISCUSSION

The SARS-CoV-2 pandemic has entered a phase in which large segments of the population have some immunity due to previous infection or vaccination. While there has been a drop in serious disease and hospitalization, variants continue to arise and spread. In the present study we investigated several topics related to vaccine efficacy in the current situation, including cross-neutralization of variants, boosting with variant S proteins, role of non-neutralizing antibody and particularly enhanced protection by IN administration of matched and mismatched vaccines. We used MVA, an attenuated vaccinia virus vector, that has been extensively used for immunological studies and is currently in clinical vaccine trials for a variety of infectious diseases including SARS-CoV-2. Before carrying out these investigations, we compared the abilities of variant SARS-CoV-2 strains to infect susceptible K18-hACE2 mice. Whereas, Washington, Beta and Delta strains were highly lethal, Omicron was less so and lower titers of the latter virus were recovered from the upper and lower respiratory tract. However, the amounts of sgRNAs in the lungs and nasal turbinates of the different strains were more similar allowing a better comparison of their replication. In addition, we constructed a panel of rMVA vaccines and a panel of rVSV pseudoviruses expressing variant S proteins. Initially we focused on the ability of rMVA-W, expressing the ancestor Wuhan S protein, to induce cross-neutralizing antibodies. In line with studies using other vaccine platforms, we found that neutralization was in the order of Wuhan > Delta > Beta > Omicron. Repeated immunizations with rMVA-W increased neutralization titers to Delta and Beta but hardly to Omicron, which has the most divergent S-protein, and none approached that to Wuhan itself. Nevertheless, vaccination with rMVA-W protected hACE2 mice against weight loss and death and reduced virus replication in the upper and lower respiratory tracts for at least 9 months. However, whereas no replication of the ancestral strain of SARS-CoV-2 was detected in the lungs by sensitive sgRNA analysis, some replication of other strains was found though significantly reduced compared to controls. Although rMVA-W elicited little anti-Omicron neutralizing antibody, there was appreciable Omicron S-binding antibody that provided partial protection when passively transferred to mice. In another study, Kaplonek and co-workers ^24^ determined that mRNA-1273 vaccine-induced antibodies maintain Fc effector functions across variants, which could explain the protection seen here. We previously reported that rMVA-W stimulated antigen-specific T cells ^12^ and the majority of the peptides in the positive pools are present in the variant S proteins. The conclusion from this phase of the study was that the mouse model mimicked clinical experience in that immunity to the ancestral SARS-CoV-2 protected against severe disease by variants but only partially prevented infection and replication.

To better understand whether the differences in neutralization and protection were mainly due to the mismatching of antibodies or to intrinsic resistance of variants to neutralization, we immunized mice with rMVAs-W, -D or -O and determined the neutralization titers to matched and mismatched pseudoviruses. In each case, neutralization of the matched pseudovirus was greater than mismatched though the difference was least between Wuhan and Delta and greatest between Omicron and the others. Nevertheless, even though rMVA-O induced antibodies that significantly neutralized rVSV-O, the titers were less than those elicited by rMVA-W for Wuhan or rMVA-D for Delta. Similar diminished cross-neutralizing antibodies were found in sera from hACE2 mice infected with sublethal doses of SARS-CoV-2 variants demonstrating that this was not a problem with the vaccines.

Another pertinent question was whether boosting mice that had been vaccinated with rMVA-W with rMVA-B, -D, or -O would increase antibodies to Wuhan S (original antigenic sin), to the variants or both. Following two vaccinations with rMVA-W, a single vaccination with rMVA-B or rMVA-D boosted the neutralization titers to Wuhan as well as to self. Although a single immunization with rMVA-O boosted neutralizing antibody to the other variants, a second vaccination with rMVA-O was required to induce neutralizing antibody to itself. Thus, two rMVA-O vaccinations were needed to raise Omicron neutralizing antibody in both naïve mice and mice that had been previously vaccinated with rMVA-W. Although not directly measured, it seems likely that in each case the first Omicron vaccination elicited Omicron-specific memory cells that were activated on the second vaccination.

By analyzing virus and sgRNAs in nasal turbinates and lungs at 2 and 4 days after SARS-CoV-2 infection of K18-hACE2 mice, we confirmed our previous data on the better protection afforded by IN compared to IM vaccination with rMVA-W. In the latter study, induction of antigen-specific IgA and higher numbers of CD8+ T cells were found in the lungs. Here we showed that IN vaccination also provided greater protection against other SARS-CoV-2 variants following immunization with matched as well as mismatched S vaccines. These studies should encourage the evaluation of nasal or aerosol vaccines to boost immunity in clinical trials.

## MATERIALS AND METHODS

### Mice

Five-to six-week-old female C57BL/6ANTac and B6.Cg-Tg(K18-hACE2)2Prlmn/J mice were obtained from Taconic Biosciences and Jackson Laboratories, respectively. Typically, 3-5 mice were housed per sterile, ventilated microisolator cage in an ABSL-2 or ABSL-3 facility.

### MVA viruses and cells

rMVA-B, rMVA-D and rMVA-O viruses were constructed as described previously ^12^. All rMVA viruses were purified by two consecutive sucrose gradients. Vero E6 cells (ATCC CRL-1586) and Vero E6 hTMPRSS2 hACE2 ^18^ were maintained in Dulbecco’s Modified Eagle Medium supplemented with 8% heat-inactivated fetal bovine serum, 2 mM L-glutamine, 10 U/ml penicillin, and 10 μg/ml streptomycin.

### Vaccination

Viruses used for vaccination were thawed, dispersed by sonication, and 10-fold serial dilutions were made in phosphate buffered saline containing 0.05% bovine serum albumin, resulting in concentrations ranging from 2×10^8^ to 2×10^4^ PFU/ml. The rMVAs in 50 μl were injected IM into each hind leg of the mouse. For IN vaccination, mice were lightly sedated with isoflurane and 50 μl of rMVAs administered.

### Infection with SARS-CoV-2

SARS-CoV-2 USA-WA1/2020 from BEI resources (Ref# NR-52281) was propagated in Vero cells (CCL81); SARS-CoV-2 Beta (RSA 1.351 501Y) from the NIAID Integrated Research Facility at Ft. Detrick; SARS-CoV-2 Delta (hCoV-19/USA/MD – HP05285/2021 VOC G/478K.V1 B.1.617.2+AY.1+AY.2) from Andrew Pekosz at Johns Hopkins University, and SARS-CoV-2 Omicron BA.1 (Ref EPI-ISL_7171744) from Vincent Munster of the NIAID Laboratory of Virology were propagated in TMPRSS2 VeroE6 cells. The clarified culture medium was titrated on Vero E6 hTMPRSS2 cells and the TCID50 was determined by the Reed-Muench method. SARS-CoV-2 were amplified and purified in a BSL-3 laboratory by Reed Johnson and Nicole Lackemeyer of the NIAID COVID Virology Core Laboratory. Aliquota consisting of 10^2^ to 5 x 10^4^ TCID_50_ of SARS-CoV-2 in 50 μl were administered IN to mice that were lightly sedated with isoflurane. After infection, the weights and morbidity/mortality status were assessed and recorded daily for up to 14 days.

### Detection of Wuhan S, Omicron S, and RBD binding IgG and antibodies by ELISA

SARS-CoV-2 (2019-nCoV) spike (S1+S2 ECD protein, Sino Biologicals), Omicron BA 1.1 spike (from Dr. Raul Cachau, NIAID) or CoV-2 Spike RBD (His-Tag, Genscript) was diluted in phosphate buffered saline (PBS) to a concentration of 1 μg/ml. MaxiSorp 96-well flat-bottom plates (Thermo Fisher) were filled with 100 μl of diluted S protein (0.1 μg/well) and incubated overnight at 4°C. After adsorption, wells were washed three times with 250 μl PBS + 0.05% Tween-20 (PBS-T, Accurate Chemical). Plates were blocked for 2 h at room temperature with 200 μl PBS-T + 5% nonfat milk and subsequently washed three times with PBS-T prior to incubation with a series of eight 4-fold dilutions of mouse sera for 1 h at room temperature. To detect S-specific IgG antibodies, plates were washed three times with PBS-T and incubated with horse radish peroxidase (HRP)-conjugated goat anti-mouse IgG (H+L) (Thermo Fisher) for 1 h at room temperature. After incubation plates were washed three times with PBS-T and 100 μl of pre-warmed SureBlue TMB substrate (SeraCare) was added to the plate for 10 min at room temperature. To stop the colorimetric reaction, 100 μl of 1N sulfuric acid was added to each well and absorbance was measured at A_450_ and A_650_ using a Synergy H1 plate reader with Gen5 analysis software (Agilent Technologies). IgG endpoint titers were determined as 4-fold above the average absorbance of those wells not containing primary antibody.

### Pseudovirus neutralization assays

BHK-21 cell lines expressing SARS-CoV-2 codon optimized spikes with truncation of the 19 C-terminal amino acids were prepared and used to generate rVSVDG–GFP-CoV2 spike pseudoviruses as previously described ^18^. For the rVSVDG pseudoviral neutralization assay, serial dilutions of heat-inactivated sera were incubated with rVSVDG pseudoviruses and anti-VSV-G I1 hybridoma supernatant (ATCC# CRL-2700) for 45 min at 37°C. The mixture was then added to VeroE6 cells expressing hTMPRSS2 and hACE2 and incubated for 20 h at 37°C. The cells were fixed in 2% paraformaldehyde and GFP measured by flow cytometry. NT50 values were calculated using Prism (Graphpad) to plot dose-response curves, normalized using the average of the no virus wells as 100% neutralization, and the average of the no serum wells as 0%. The limit of detection (LOD) of 25 was determined by taking 1.96 standard deviation of the mean titer of the control MVA samples.

### Quantitation of infectious SARS-CoV-2

Lungs, brains, and nasal turbinates were homogenized, cleared of debris by centrifugation at 3800xg for 10 min and serial 10-fold dilutions were applied in quadruplicate to Vero E6 hTMPRSS2 cells in DMEM+Glutamax (ThermoFisher) supplemented with 2% heat-inactivated FBS and 1% Antibiotic-Antimycotic in 96-well microtiter plates. After 72 h, the plates were stained with crystal violet and the Reed-Muench method was used to determine the concentration at which 50%of the cells displayed a cytopathic effect (TCID_50_).

### Quantitation of SARS-CoV-2 sgRNAs

RNA was extracted from homogenates of lungs and turbinates using Trizol; contaminating DNA was removed and RNA was reverse-transcribed. SARS-CoV-2 sgS and sgN transcripts and 18S rRNA were quantified by ddPCR with specific primers using an automated droplet generator and droplet reader (BioRad).

### Safety and Ethics

All experiments and procedures involving mice were approved under protocol LVD29E by the NIAID Animal Care and Use Committee according to standards set forth in the NIH guidelines, Animal Welfare Act, and US Federal Law. Euthanasia was carried out using carbon dioxide inhalation in accordance with the American Veterinary Medical Association Guidelines for Euthanasia of Animals (2013 Report of the AVMA Panel of Euthanasia). Experiments with SARS-CoV-2 were carried out under BSL-3 containment.

## Data Availability

All data is included in the manuscript and supporting information.

## ACKNOWLEDGEMENTS

We thank Reed Johnson and Nicole Lackemeyer of the NIAID COVID Virology Core Laboratory and Vincent Munster of the NIAID Laboratory of Virology for stocks of SARS-CoV-2. The technical staff of the NIAID Comprehensive Medical Branch provided excellent animal care. The work was supported by the Division of Intramural Research of NIAID.

## AUTHOR CONTRIBUTIONS

B.M. and P.E. designed experiments, C.A.C. and J.L.A. carried out experiments, B.M. wrote the paper, C.A.C. prepared the figures, and all authors edited the final manuscript.

## COMPETING INTEREST STATEMENT

The authors declare no competing interest.

## FIGURE LEGENDS

**Fig. S1.**
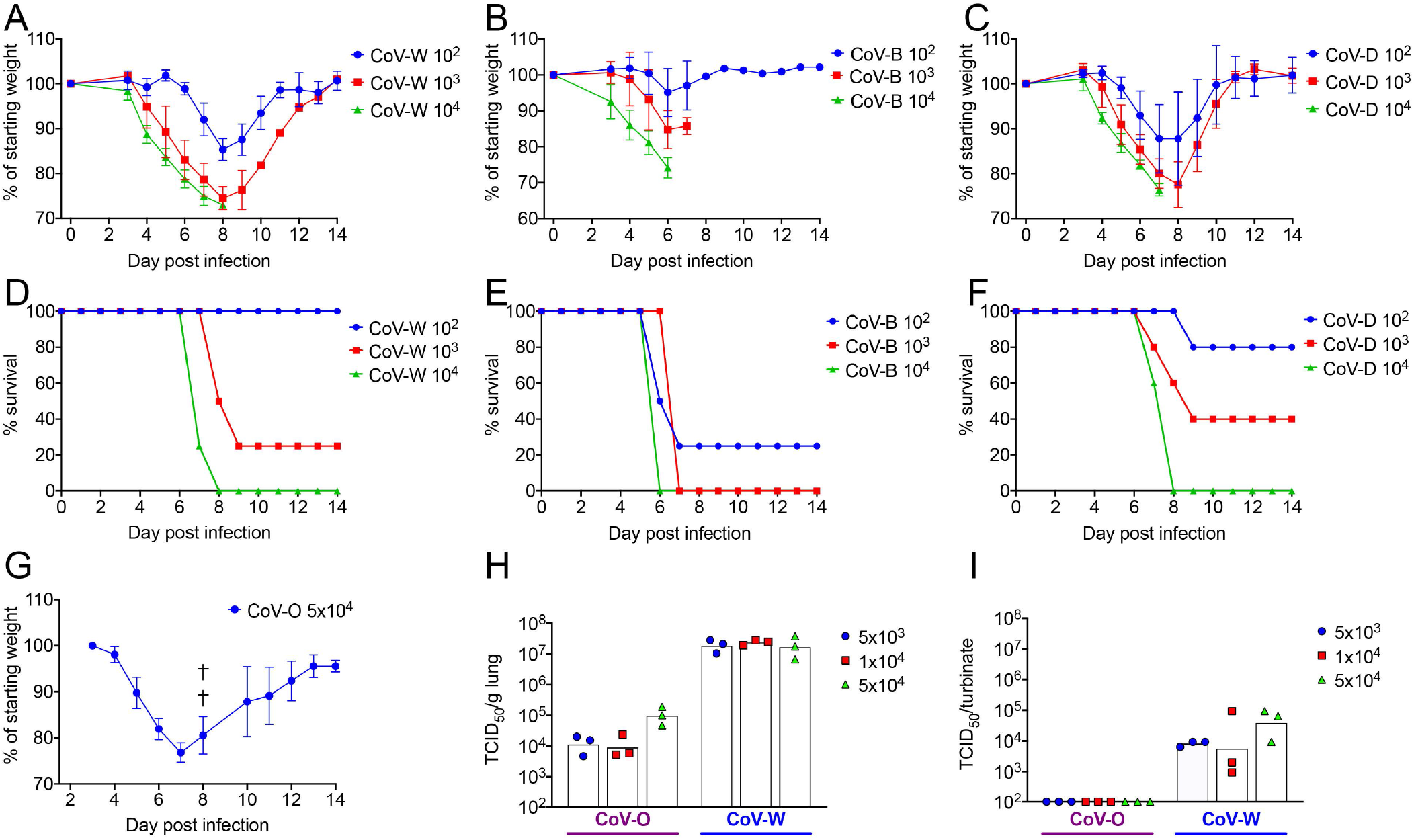
Relative virulence of SAR-CoV-2 variants. (**A -F**) K18-hACE2 mice (n=3 per group) were infected IN with 10^2^ to 10^4^ TCID50 of indicated SARS-CoV-2 strains Washington (CoV-W, Beta (CoV-B), Delta (CoV-D) and weight loss and survival plotted. (G) K18-hACE2 mice (n=5) were infected IN with 5 x 104 TCID50 of CoV-O and weight loss plotted. †, death. (**H, I**) K18-hACE2 mice (n=3 per group) were infected IN with 5 x 10^3^ to 5 x 10^4^ TCID_50_ of CoV-W or CoV-O and virus titers in the lungs and nasal turbinates determined on day 2.

**Fig. S2.**
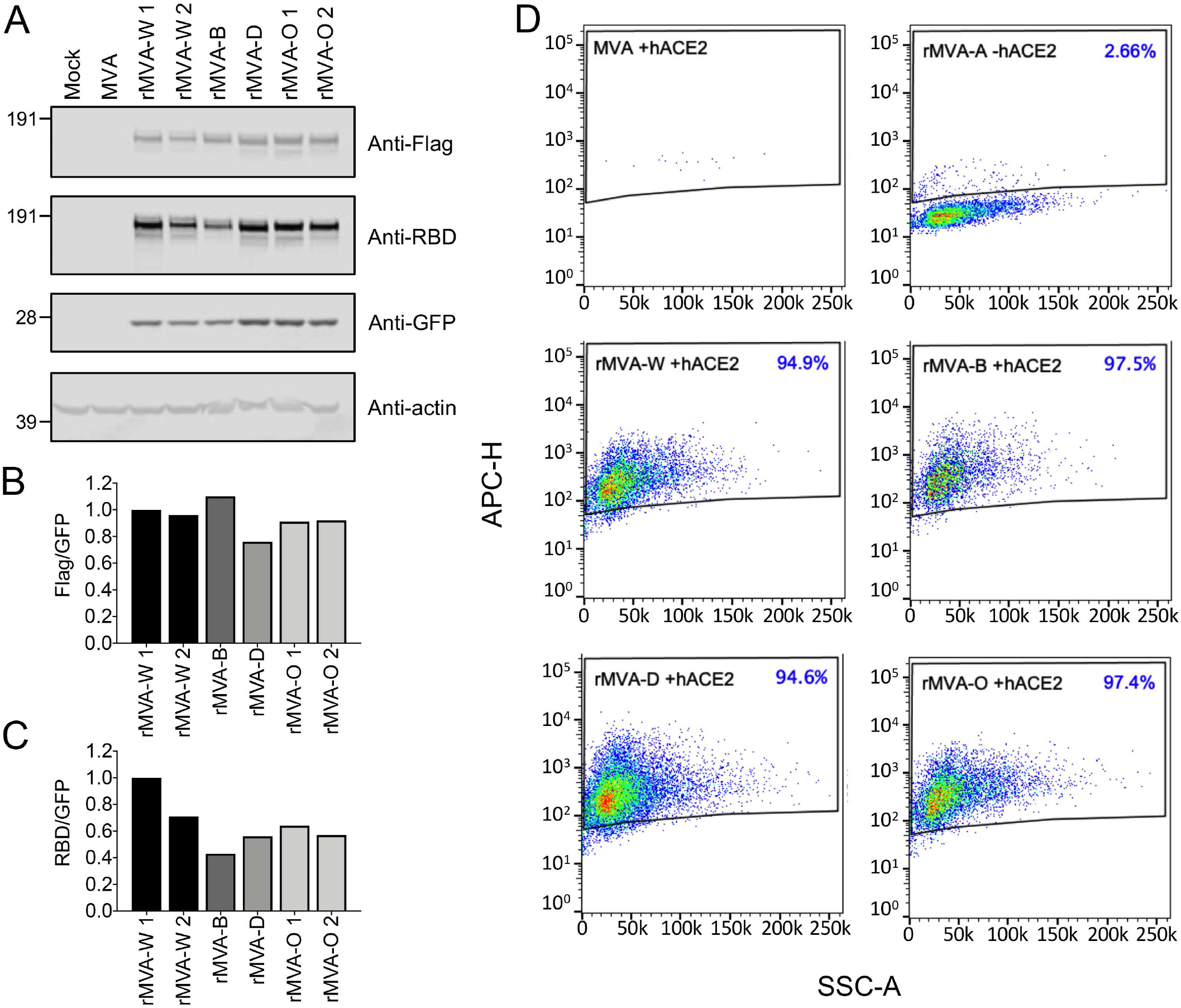
Expression of S by variant rMVAs. **(A)** Cells were mock-infected or infected with rMVA-W, -B, -D, or -O and Western blots probed with mouse anti-Flag (Millipore Sigma), rabbit anti-Wuhan S-RBD (Sino Biologicals), rabbit anti-GFP (Thermofisher), mouse anti-actin (Santa Cruz), donkey anti-mouse IRDye 680RD (LiCor) and donkey anti-rabbit IRDye800CW (LiCor). The positions of size markers with mass in kDa are shown on the left. **(B)** Ratios of intensities of the bands probed with anti-Flag and anti-GFP plotted. **(C)** Ratios of intensities of bands probed with anti-RBD and anti-GFP plotted. **(D)** Cells were infected with MVA, rMVA-A, -W, -B, -D and -O, fixed without permeabilization and incubated with biotinylated his-tag hACE2 (Sino Biologicals) followed by APC streptavidin (BD Pharmagin). Stained cells were analyzed by flow cytometry. Cells were first gated for GFP fluorescence and then for APC. Percent of GFP+ cells that stained with hACE2 are indicated.

**Fig. S3.**
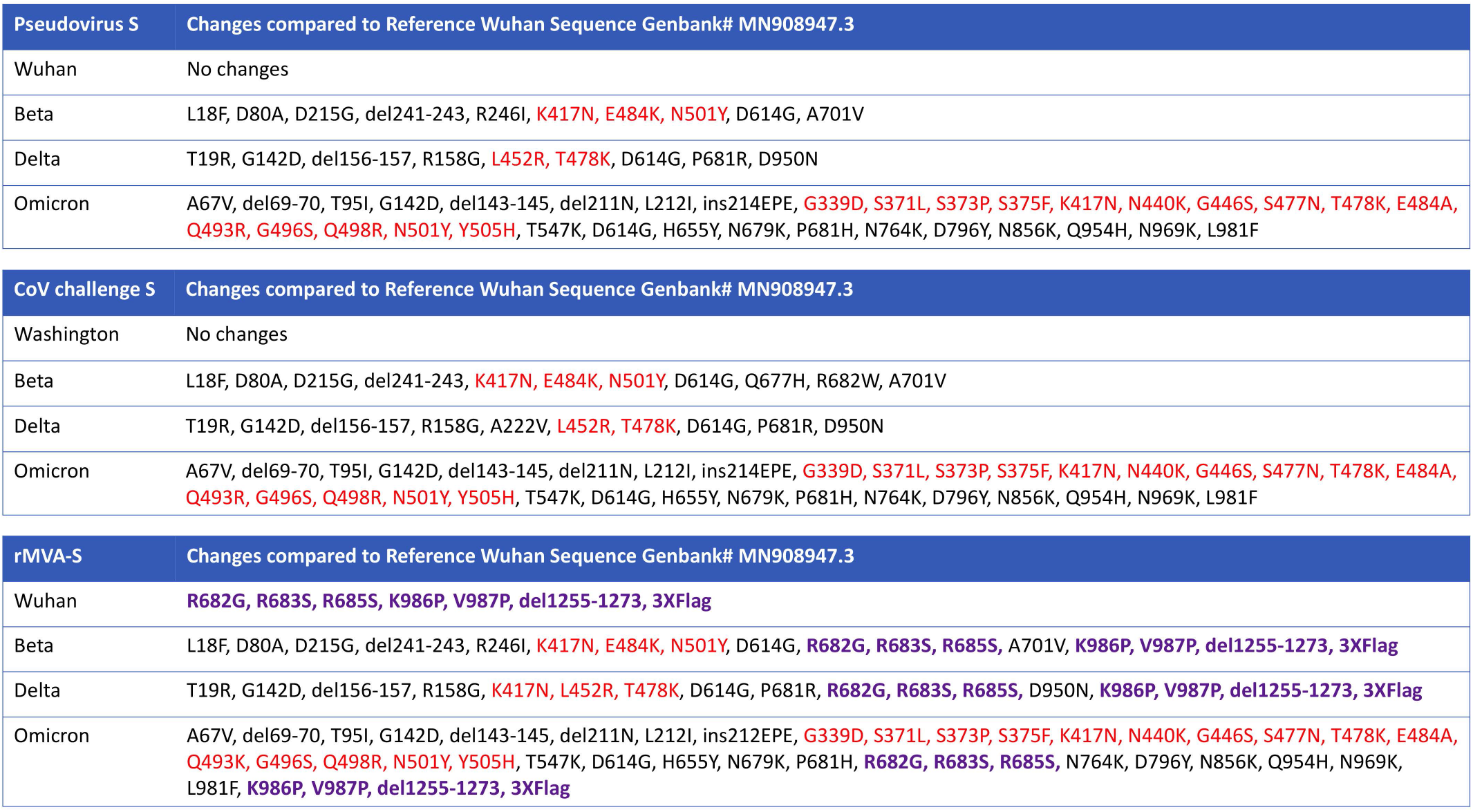
Comparison of sequences of S in rMVAs, rVSVs and SARS-CoV-2 variants. All sequences are compared to Wuhan Genbank# MN908947.3 and only differences are listed. Amino acids in red are in RBD; amino acids in purple represent modifications in rMVAs for stability of the prefusion form of S, prevent furin cleavage and endoplasmic retrieval, and add a 3 x Flag tag as previously described for Wuhan ^12^. Abbreviation: del, deletion.

## Notes

### Competing Interest Statement

The authors have declared no competing interest.

## REFERENCES

1 Polack, F. P. et al. Safety and Efficacy of the BNT162b2 mRNA Covid-19 Vaccine. N Engl J Med 383, 2603–2615, doi:10.1056/NEJMoa2034577 (2020).

2 Baden, L. R. et al. Efficacy and Safety of the mRNA-1273 SARS-CoV-2 Vaccine. N Engl J Med 384, 403–416, doi:10.1056/NEJMoa2035389 (2021).

3 Sadoff, J. et al. Safety and Efficacy of Single-Dose Ad26.COV2.S Vaccine against Covid-19. N Engl J Med 384, 2187–2201, doi:10.1056/NEJMoa2101544 (2021).

4 Falsey, A. R. et al. Phase 3 Safety and Efficacy of AZD1222 (ChAdOx1 nCoV-19) Covid-19 Vaccine. N Engl J Med 385, 2348–2360, doi:10.1056/NEJMoa2105290 (2021).

5 Robson, F. et al. Coronavirus RNA Proofreading: Molecular Basis and Therapeutic Targeting. Mol Cell 80, 1136–1138, doi:10.1016/j.molcel.2020.11.048 (2020).

6 Harvey, W. T. et al. SARS-CoV-2 variants, spike mutations and immune escape. Nat Rev Microbiol 19, 409–424, doi:10.1038/s41579-021-00573-0 (2021).

7 Riemersma, K. K. et al. Shedding of infectious SARS-CoV-2 despite vaccination. PLoS Pathog 18, e1010876, doi:10.1371/journal.ppat.1010876 (2022).

8 Chalkias, S. et al. A Bivalent Omicron-Containing Booster Vaccine against Covid-19. N Engl J Med 387, 1279–1291, doi:10.1056/NEJMoa2208343 (2022).

9 Moss, B. Genetically engineered poxviruses for recombinant gene expression, vaccination, and safety. Proc. Natl. Acad. Sci. USA 93, 11341–11348 (1996).

10 Volz, A. & Sutter, G. in Advances in Virus Research, Vol 97 Vol. 97 Advances in Virus Research (eds M. Kielian, T. C. Mettenleiter, & M. J. Roossinck) 187–243 (2017).

11 Chiuppesi, F. et al. Safety and immunogenicity of a synthetic multiantigen modified vaccinia virus Ankara-based COVID-19 vaccine (COH04S1): an open-label and randomised, phase 1 trial. Lancet Microbe 3, e252–e264, doi:10.1016/S2666-5247(22)00027-1 (2022).

12 Liu, R. K. et al. One or two injections of MVA-vectored vaccine shields hACE2 transgenic mice from SARS-CoV-2 upper and lower respiratory tract infection. Proceedings of the National Academy of Sciences of the United States of America 118, doi:10.1073/pnas.2026785118 (2021).

13 Routhu, N. K. et al. A modified vaccinia Ankara vector-based vaccine protects macaques from SARS-CoV-2 infection, immune pathology, and dysfunction in the lungs. Immunity 54, 542-+, doi:10.1016/j.immuni.2021.02.001 (2021).

14 Tscherne, A. et al. Immunogenicity and efficacy of the COVID-19 candidate vector vaccine MVA-SARS-2-S in preclinical vaccination. Proc Natl Acad Sci U S A 118, doi:10.1073/pnas.2026207118 (2021).

15 Garcia-Arriaza, J. et al. COVID-19 vaccine candidates based on modified vaccinia virus Ankara expressing the SARS-CoV-2 spike induce robust T-and B-cell immune responses and full efficacy in mice. J Virol, doi:10.1128/JVI.02260-20 (2021).

16 Meseda, C. A. et al. MVA vector expression of SARS-CoV-2 spike protein and protection of adult Syrian hamsters against SARS-CoV-2 challenge. Npj Vaccines 6, doi: 10.1038/s41541-021-00410-8 (2021).

17 Bosnjak, B. et al. Intranasal Delivery of MVA Vector Vaccine Induces Effective Pulmonary Immunity Against SARS-CoV-2 in Rodents. Frontiers in Immunology 12, doi:10.3389/fimmu.2021.772240 (2021).

18 Americo, J. L., Cotter, C. A., Earl, P. L., Liu, R. & Moss, B. Intranasal inoculation of an MVA-based vaccine induces IgA and protects the respiratory tract of hACE2 mice from SARS-CoV-2 infection. Proc Natl Acad Sci U S A 119, e2202069119, doi:10.1073/pnas.2202069119 (2022).

19 Perez, P. et al. Intranasal administration of a single dose of MVA-based vaccine candidates against COVID-19 induced local and systemic immune responses and protects mice from a lethal SARS-CoV-2 infection. Frontiers in Immunology 13, doi:10.3389/fimmu.2022.995235 (2022).

20 Halfmann, P. J. et al. SARS-CoV-2 Omicron virus causes attenuated disease in mice and hamsters. Nature, doi:10.1038/s41586-022-04441-6 (2022).

21 Suryawanshi, R. K. et al. Limited cross-variant immunity from SARS-CoV-2 Omicron without vaccination. Nature 607, 351–355, doi:10.1038/s41586-022-04865-0 (2022).

22 Speranza, E. et al. Single-cell RNA sequencing reveals SARS-CoV-2 infection dynamics in lungs of African green monkeys. Sci Transl Med 13, doi:10.1126/scitranslmed.abe8146 (2021).

23 Kim, D. et al. The Architecture of SARS-CoV-2 Transcriptome. Cell 181, 914–921 e910, doi:10.1016/j.cell.2020.04.011 (2020).

24 Kaplonek, P. et al. mRNA-1273 vaccine-induced antibodies maintain Fc effector functions across SARS-CoV-2 variants of concern. Immunity 55, 355–365 e354, doi:10.1016/j.immuni.2022.01.001 (2022).

